# Variations in clustering of multielectrode local field potentials in the motor cortex of macaque monkeys during a reach-and-grasp task

**DOI:** 10.1101/2024.03.20.585852

**Authors:** Florian Chambellant, Ali Falaki, Ian Moreau-Debord, Robert French, Eleonore Serrano, Stephane Quessy, Numa Dancause, Elizabeth Thomas

## Abstract

There is experimental evidence of varying correlation among the elements of the neuromuscular system over the course of the reach-and-grasp task. Several neuromuscular disorders are accompanied by anomalies in muscular coupling during the task. The aim of this study was to investigate if modifications in correlations and clustering can be detected in the Local Field Potential (LFP) recordings of the motor cortex during the task. To this end, we analyzed the LFP recordings from a previously published study on monkeys which performed a reach-and-grasp task for targets with a vertical or horizontal orientation. LFP signals were recorded from the motor and premotor cortex of macaque monkeys as they performed the task. We found very robust changes in the correlations of the multielectrode LFP recordings which corresponded to task epochs. Mean LFP correlation increased significantly during reaching and then decreased during grasp. This pattern was very robust for both left and right arm reaches irrespective of target orientation. A hierarchical cluster analysis supported the same conclusion – a decreased number of clusters during reach followed by an increase for grasp. A sliding window computation of the number of clusters was performed to probe the predictive capacities of these LFP clusters for upcoming task events. For a very high percentage of trials (95.3%), there was a downturn in cluster number following the Pellet Drop (GO signal) which reached a minimum shortly preceding the Start of grasp, hence indicating that cluster analyses of LFP signals could provide online indications of the Start of grasp.

## Introduction

The optimal functioning of muscular activity requires that they couple and uncouple to the right extent in a timely manner. The breakdown of this coordination very clearly leads to dysfunction of the upper limbs for our daily tasks. An example of this is the excessive muscular co-activation patterns seen during the early stages following stroke. During this state, muscles show poor ability to contract in isolation giving rise to either extensor or flexor synergies (Baak et al., 2015; Brunnstrom, 1970; Twitchell, 1951). In hand functioning, the power grip which requires less finger individuation is favored in these conditions (Li et al., 2003; Xu et al., 2015). At the other end of the continuum are situations of poor interjoint coordination for some movement disorders (Astill & Utley, 2008; Levin, 1996).

Given the importance of correlations in muscular activity, it is natural to expect that we might find this reflected in the neural command structure. Some research groups have found that neural stimulation is able to elicit movements with low dimensional stereotypical patterns. For example, Transcranial Magnetic Stimulation (TMS) in the primary motor cortex can elicit hand movements of low dimensionality with a modular architecture (Bufalari et al., 2010; Gentner & Classen, 2006; Schwenkreis et al., 2007). While these examples of correlation remain expressed at the muscular level, more direct evidence of such clustering at the neural level are also available, e.g. Overduin et al. (Overduin et al., 2015) demonstrated the presence of low dimensional neural activation patterns during reach-and-grasp in multiunit recordings of the motor cortex of monkeys. Leo et al. (Leo et al., 2016) predicted hand postures from corresponding low dimensional representations of fMRi data from the motor cortex.

In the present study we continue with the effort to characterize this organization of neural commands in association with the reach-and-grasp movements. In contrast to the previous studies on low dimensional activities in the motor cortex using neuronal spiking activity (Overduin et al., 2012, 2015) and fMRI (Leo et al., 2016), we used local field potentials (LFPs). As a low frequency, extracellular signal, it is known to be relatively robust and less subject to factors such as electrode position (Bédard et al., 2004; Scherberger, 2009). We aimed to investigate the correlations between multielectrode LFP activities in the motor cortex. In particular we wanted to know if they would change over the course of task execution. To this end, we analyzed signals collected from multielectrode arrays implanted in the primary motor cortex (M1) of the right hemisphere as well as in the dorsal and ventral premotor cortex (PMd and PMv, respectively) of both hemispheres of macaque monkeys of a previous investigation (Moreau-Debord et al., 2021).

Few studies have reported on how the correlations of LFP activity in the motor cortex evolves as a function of reach-and-grasp task epochs. Spinks et al. (Spinks et al., 2008) conducted a study of mean correlation within and between M1 and PMv without examining the evolution of this correlation as movement unfolds. Scherberger et al. (Scherberger et al., 2005) examined correlations between field recordings and single unit spiking activity. Our study moves the field ahead by asking if and how clustering properties between LFP electrodes change during the reach-and-grasp task.

To achieve our aims we used a Pearson correlation analysis followed by a hierarchical clustering algorithm to examine in more detail how the correlations between the electrodes within the motor cortex evolved over the course of the reach-and-grasp task. Hierarchical clustering has often been used to examine both physical and functional clustering in the field of Neuroscience (Boly et al., 2012; Honig et al., 2021; Liu et al., 2012; Merchant & Georgopoulos, 2006). A more pertinent example is a very recent study by Quarta et al. (Quarta et al., 2022) where hierarchical clustering on calcium imaging signals was used to investigate neocortical activity during a reach-and-grasp task.

## Methods

### Experimental model and surgery

The data were recorded in two female rhesus macaque (Macaca mulatta) monkeys - Monkey M (5.5 kg) and Monkey S (5.7 kg). Details on the surgical procedures and the behavioral task have been previously published (Moreau-Debord et al., 2021). Briefly, animals underwent craniotomies and durectomies to expose M1 and the lateral premotor cortex in both hemispheres. Multielectrodes arrays were implanted in the PMv and PMd of both hemispheres, as well as the M1 of the right hemisphere.

LFP data were collected and sampled at a 2035 Hz and neuronal data spiking data at 24414 Hz with a Tucker-Davis Technologies acquisition system. The recordings were filtered between 0-500 Hz to extract the LFPs while spiking data were band-pass filtered between 100-5000 Hz and sorted offline using Plexon Offline Sorter (Plexon) to obtain spikes times for each electrode.

As the acquisition device was only able to record 256 channels simultaneously, a different subset of electrodes were selected for each recording session (total number of implanted electrodes: 370 electrodes in Monkey M and 448 electrodes in Monkey S). Here, we present results from two sessions recorded in monkey M and two sessions in monkey S during which neuronal recordings were obtained from most of the implanted brain areas.

### Behavioral task

The monkeys sat on a custom-made primate chair placed in front of a food pellet dispenser. The chair was equipped with an opening for the mouth and two removable panels, one at each side of the chair, allowing the monkey to use one hand to withdraw the food pellets and eat them. Either the left or the right opening for the arm (depending on the trial block) was available for the monkey to pass its hand through and retrieve the food pellet dropped in a well behind a slot (1.3 cm × 5.5 cm) located about 10 cm below the shoulder height and 20 cm from the monkeys. Depending on the trial block, the slot had either a horizontal or vertical orientation hence forcing the animal to appropriately supinate or pronate the hand and obtain the pellet using a precision grip with the thumb and index (See Figure 1a for example of vertical grasp).

**Figure 1.**
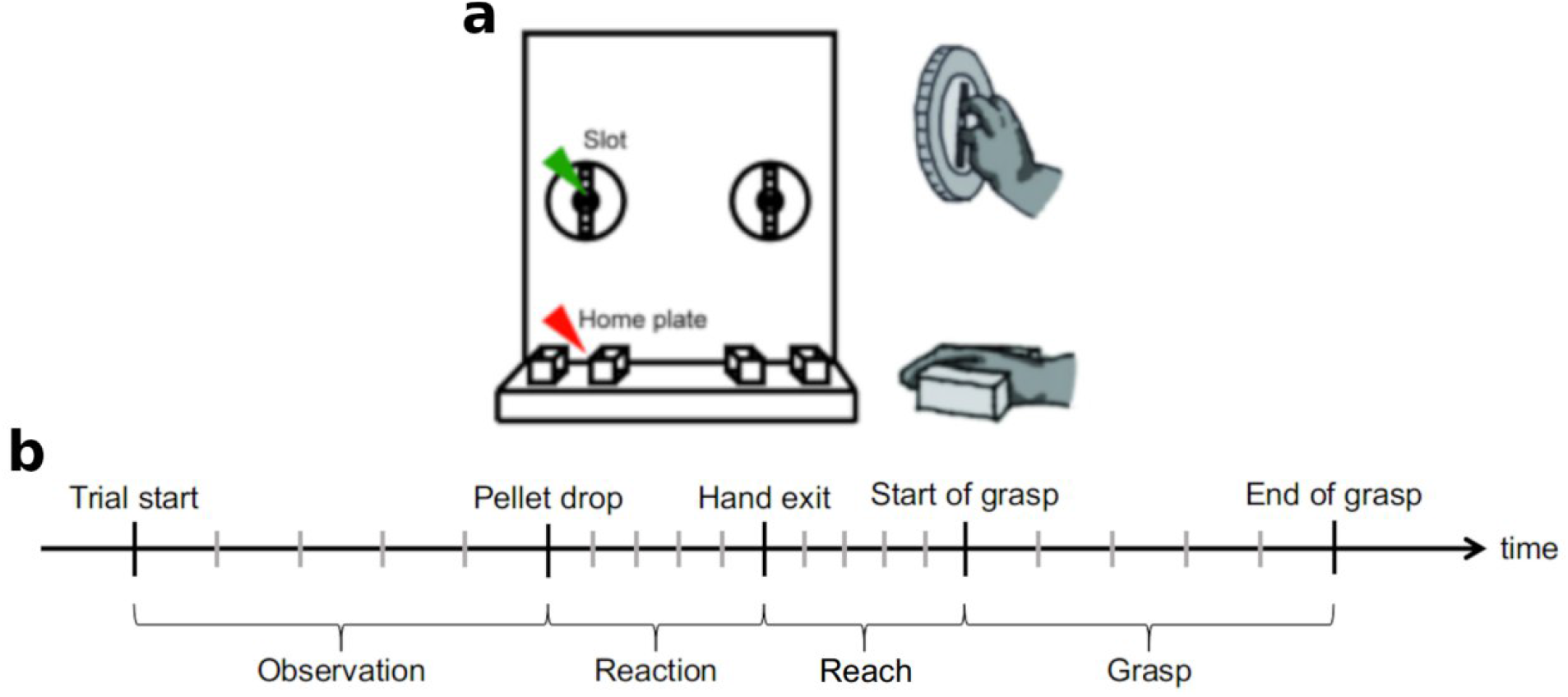
a) Experimental apparatus (adapted from (Moreau-Debord et al. 2021)). b) Trial segmentation and events. More details on the manner in which each epoch was defined can be found in the Methods section.

Each trial began after the animal placed its hand on the home plate located 15 cm below the pellet slot. The presence of the hand was detected by an infrared laser sensor embedded in the home plate and marked as ‘Trial start’. After a random delay ranging from 800 ms to 2 s, a pellet was dropped into the well (Pellet drop) using a pellet dispenser (80209 Pellet Dispenser, Campden Instrument). The sound resulting from the pellet drop served as a GO cue for the monkey. Following the pellet drop, the animal had 2 s to remove its hand from the home plate (Hand exit) and Reach towards the pellet. The time when the animal entered its hand in the slot to grasp the pellet (Start of grasp) and when the hand was removed from the slot (End of grasp) were detected by a second infrared laser sensor embedded at the entrance of the slot. The monkey brought the pellet to its mouth and placed its hand back on the home plate to initiate the next trial with an intertrial interval of 3 s. The animals repeated the task 25 times for each hand and slot orientation (block randomized). See Figure 1b for an indicative timeline of the task.

### Preprocessing

Custom scripts written in Matlab (The MathWorks Inc., 2021) and the Fieldtrip toolbox (Oostenveld et al., 2010) were used to process neural data. Prior to performing the correlation and clustering analyses, neural data were preprocessed to remove bad trials as well as potential noises and artifacts. Trials for which the animal’s Reaction time was inferior to 200 ms (the monkey successfully predicted the pellet drop), the Reach duration was above 350 ms or the grasping lasted less than 200 ms (the monkey was unsuccessful in retrieving the pellet in a single try) were discarded. This represented 5.2% of the trials. Following the rejection of bad trials, the LFP data were cleaned using a low-pass Butterworth filter (200 Hz, 6^th^ order) followed by the removal of 60 Hz noise using notch Butterworth filter with band stop limits of 59 and and 61Hz. Irregular bursts and low frequency irregular oscillations were removed using independent component analysis (ICA) (Bell & Sejnowski, 1995; Whitmore & Lin, 2016). Since we collected LFP data in blocks of 32 electrodes using Omnetics connectors, the ICA procedure was applied to the data collected by each individual connector to reduce spreading signal cancellation. When the ICA procedure failed i.e. more than 10 components contained noise, data from the entire connector was discarded. Following these procedures, electrodes with outlier LFP activity i.e. greater than the mean ± 2 std were also removed. Further preparation of the LFP data consisted of removing effects of neuronal spiking activity. Recoded information on the spike timings were used to replace the data of an electrode, corresponded to the spiking activity, from 1 ms before to 2 ms after the spike time by linear interpolation between these two points (Waldert et al., 2013). The number of trials and electrodes for each area for the 2 sessions of the 2 monkeys can be seen in Table 1.

**Table 1.**
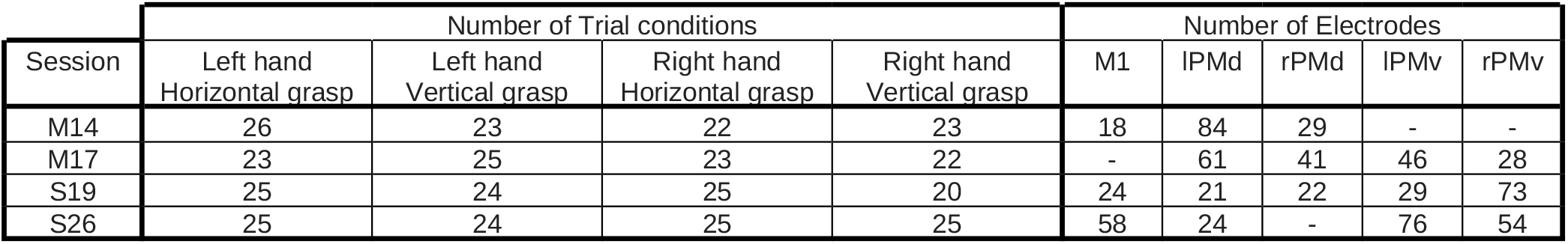
Summary of the different analyzed sessions in the study.

| Session | Number of Trial conditions |  |  |  | Number of Electrodes |  |  |  |  |
| --- | --- | --- | --- | --- | --- | --- | --- | --- | --- |
|  | Left hand<br>Horizontal grasp | Left hand<br>Vertical grasp | Right hand<br>Horizontal grasp | Right hand<br>Vertical grasp | M1 | IPMd | rPMd | IPMv | rPMv |
| M14 | 26 | 23 | 22 | 23 | 18 | 84 | 29 | - | - |
| M17 | 23 | 25 | 23 | 22 | - | 61 | 41 | 46 | 28 |
| S19 | 25 | 24 | 25 | 20 | 24 | 21 | 22 | 29 | 73 |
| S26 | 25 | 24 | 25 | 25 | 58 | 24 | - | 76 | 54 |

Finally, the power spectrograms of the cleaned data were obtained using the single taper method (400 ms sliding window with 10 ms time steps) and were normalized by the average activity between 600 ms and 200 ms before Pellet drop (decibel normalization) (Menzer et al., 2014; Waldert et al., 2015) (Figure 2).

**Figure 2.**
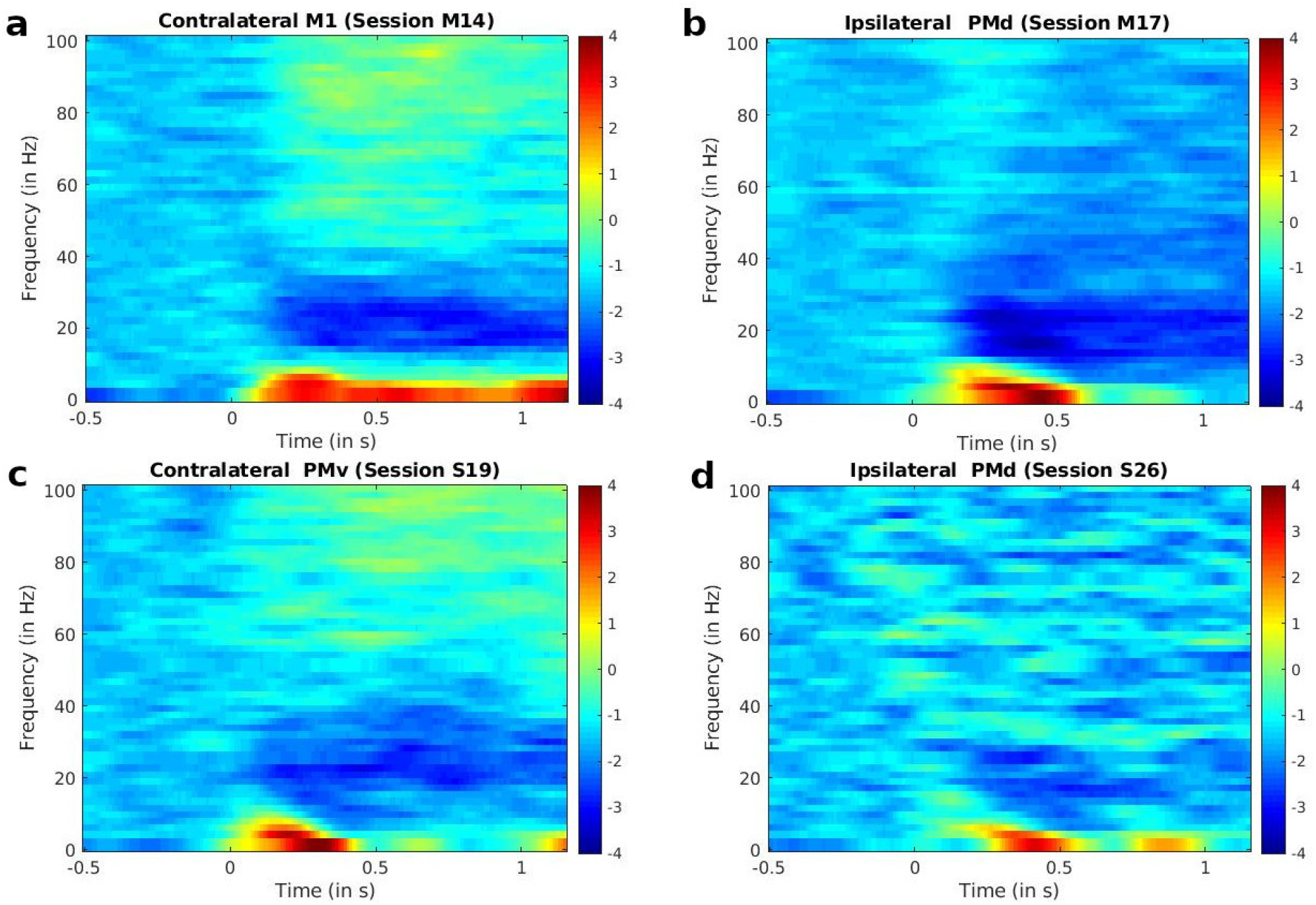
Examples of spectrograms of the LFP activity, during different sessions, aligned at Pellet drop (Time = 0) (Pellet drop served as the GO cue) averaged across all trials and electrodes for the indicated movement type (ipsilateral or contralateral) and brain area of the session.

It can be observed in Figure 2 that the highest spectral powers were observed in the delta band (approx. 0-4Hz).

### Correlation

The similarity in the spectral power of the LFP activities over the course of the reach-and-grasp task was first analyzed using a correlation analysis. Each trial was divided into 4 phases based on the events of the trial, i.e. Observation, Reaction, Reach, and Grasp (Figure 1a and b). In all cases, correlations were computed across all electrodes irrespective of the area of the motor cortex in which they were implanted. For each trial, average correlations was obtained from the absolute values of the pairwise correlations between all electrodes of the trial. In cases where we were displaying the correlations from a session, the correlation values were averaged across trials. While in Figure 4a, we display the correlation at all the frequencies, the other analyses of the paper were focused on the delta frequency band, so that the final correlation values analyzed were an average of correlations across the delta band.

### Hierarchical clustering

A finer analysis of the proximity of activities between the LFP electrodes was obtained using a cluster analysis. In this method, a distance measure is used to cluster together similar data points. The idea of hierarchical clustering (Hastie et al., 2009; Ward, 1963) is to create a hierarchy of similarity between data points. This hierarchy is usually presented as a dendrogram, with each branch containing data points which are close. The distance between any two clusters is represented by the length of the line holding apart the clusters (for an example, see Figure 3b).

**Figure 3.**
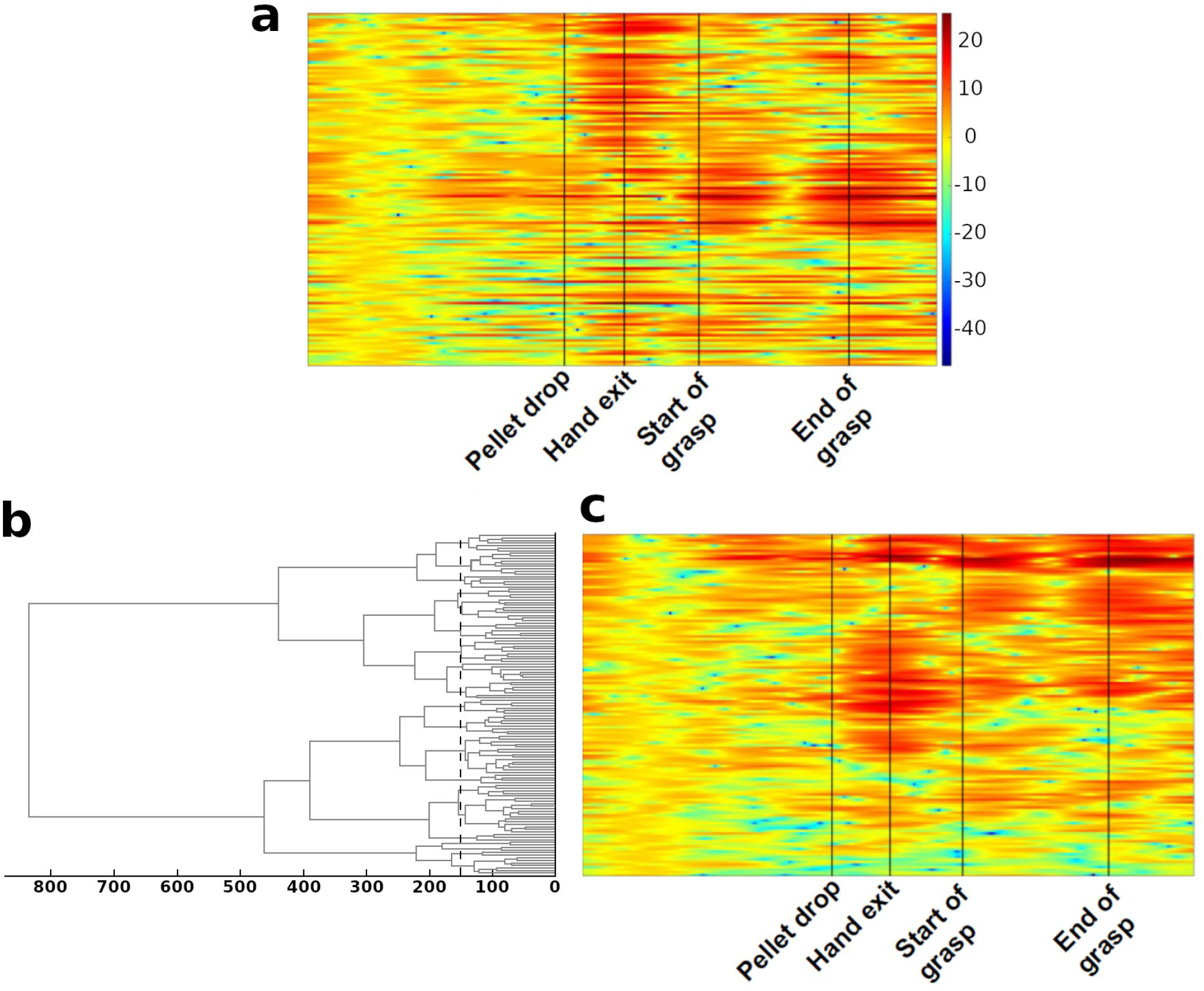
Illustration of hierarchical clustering technique. a) LFP spectral power at 2.5Hz (scale on the right) for all the electrodes from a single trial. Each horizontal line corresponds to the LFP spectral power of one recorded electrode for the entire trial. The trial epochs are indicated by vertical black lines b) Dendrogram obtained by using hierarchical clustering on the data of a). The dashed black vertical line indicates a distance of 150 (a.u.). This line was used as the point at which cluster numbers were counted. c) Data of spectrogram in a) reorganized to follow the dendrogram presented in b. The clustering algorithms was applied to the entire trial and led to the clustering of information from electrodes with correlated spectral power.

In our case, the distance measure used to evaluate the separation between two LFP electrodes was correlation. The first step of our hierarchical clustering was to compute a matrix of absolute correlations between each pair of electrodes. The LFP spectral powers of the two most correlated electrode pairs were then ‘fused’ together and a new correlation matrix was obtained with the ‘fused’ data. This process was repeated until all the points were fused together. Following this, a distance criterion was used to count the number of clusters at a fixed point. Each potential cluster was composed of a branch and all its ‘leaves’ at this distance (Figure 3b). The cutoff distance for estimating cluster number was fixed by observing the point at which there was a sharp drop in cluster number (this is similar to the elbow method used with PCA analyses to count the number of components). This distance criterion was set during the Observation phase. This criterion was then used to compute cluster numbers for each trial and sub-period of each session. As we were more interested in the changes of cluster number with the progression of the reach-and-grasp task rather than the actual number of clusters itself, the measure that was finally used to follow the similarity in LFP activity was normalized cluster number i.e. the cluster number at Pellet drop was set to 1.

The hierarchical clustering was applied in two ways. In one case, the entire trial was divided into sub-periods and the cluster numbers were computed for each sub-period (see Figure 1b). In the second case, the cluster numbers were computed using a sliding window, hence giving us an instantaneous cluster number. In both cases, the correlation matrix of the LFP signals in the delta band during the time period of interest was computed followed by the application of the hierarchical clustering method to the matrix.

### Statistical tests

Repeated measures ANOVA and Student’s t-test were used as the main statistical tests. In the case of using the ANOVA, post-hoc analysis was conducted using Tukey’s HSD test. The data were checked for assumptions of normality and sphericity and were transformed accordingly to satisfy statistical assumptions of normality. If the data did not respect the assumption of sphericity, p-values were corrected using the Greenhouse-Geisser correction and the Holm’s test was then used for post-hoc analyses. For all statistical tests, th < 0.05 was considered as the statistically significant value.

## Results

In the sections below we present the results from the correlation and clustering analyses which were performed on the LFP data during the reach-and-grasp task by two monkeys during their two sessions. The computation of correlations and clusters was done across several motor areas without taking into account the specific area of the motor cortex in which the electrodes were found. This is in keeping with the idea that correlation and cluster formation in the brain would reflect what has been observed in human behavioral studies i.e. varying degrees of coupling between the shoulder, upper arm, lower arm or hand, during reaching and grasping movements (d’Avella et al., 2008; D’avella & Lacquaniti, 2013; Jeannerod, 1984; Wallace et al., 1990). The analyses of the study were focused on the delta band as we found the largest modulations of spectral power in this band following pellet drop as seen in the spectrograms of Figure 2. Others have also reported the link between activity in the delta band of the motor cortex and arm movement (Milekovic et al., 2015; Rickert et al., 2005; Waldert et al., 2015; Zhuang et al., 2010).

### Pairwise correlation of LFP spectral power in the delta band

The correlation analyses provided us with a picture of the evolution of similarities between the recorded multielectrode LFPs over the course of the reach-and-grasp task. The correlation analysis were first done by looking at the spectral power for all frequencies of one trial of monkey S (Figure 4a). We then moved on from there to narrowing the investigation down to the delta frequency (frequency with the highest spectral power) for all the trials in one session for the same monkey S (Figure 4b). The session included different pointing conditions – the arm used and target orientation. Following this, correlation analyses were conducted separately for the different pointing conditions. This was done in order to see how generalized the patterns of correlations transitions were. Finally in Figure 4c, we computed the averages for all the trials for all task conditions of both sessions for both monkeys in order to test statistical significance. Below, we provide more details on each figure.

**Figure 4.**
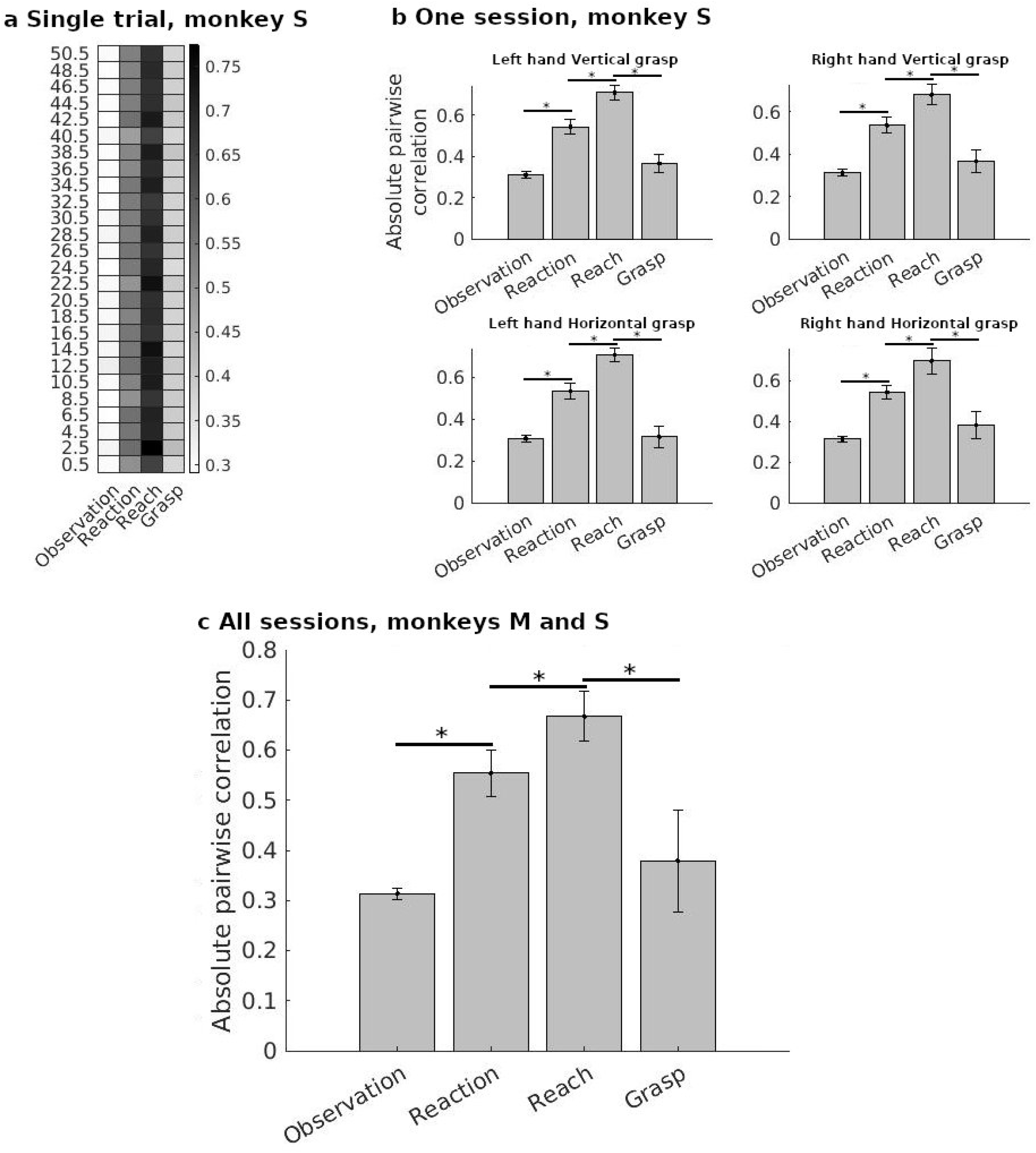
a) Heatmap of the average absolute pairwise correlation of LFP spectral power between electrodes at different frequencies during each phase of the movement for one trial of monkey S. b) Average absolute pairwise correlation of spectral power for the delta band for all the trials in one session for monkey S under different trial conditions. Error bars indicate the variance between trials. c) Average of the absolute pairwise correlation in the delta band for all trials and sessions performed by the two monkeys (two sessions per monkey). Note that this included different pointing conditions. Error bars indicate the variance between trials.

Figure 4a displays the correlation values for one trial performed by monkey S, during which the left arm was used to perform a horizontal grasp movement. The values shown are the averages of the absolute values of pairwise correlations in the LFP spectral power at each frequency between all the recorded electrodes (see Methods). The LFP correlations between the different task epochs were significantly different for this representative trial (Repeated measures ANOVA, th(1.735,671.303) = 2672.209,th < 0.001, Greenhouse-Geisser correction for sphericity). More specifically, as compared to the Observation period, there was an increase in the average absolute pairwise correlation during the Reaction time with further increase during the Reach phase. Finally, the average absolute pairwise correlation decreased during the Grasping phase. The correlation changes during all these task epoch transitions were found to be significant using post-hoc analyses (Holm’s test, th < 0.001 for all transitions).

The results displayed in Figure 4a were obtained from one trial of one type of reach-and-grasp, namely, a left arm reach for a horizontal grasp. However, the monkeys in this study also performed trials with right arm reaches and vertical grasps. In Figure 4b we examine the profile of correlation transitions for the same monkey S under these different task conditions. In contrast to Figure 4a, in Figure 4b we focused on the spectral power of the delta band. We first computed the pairwise correlations between the electrodes of single trials, before averaging the correlations over the pertinent trials. Each figure for the different pointing types in Figure 4b, is the average of all the trials with the same task conditions in that session. Similar profiles of LFP correlation increases and decreases were seen in the different types of reach-and-grasp task conditions for monkey S.

A final test was done to examine the statistical significance of these transitions for both monkeys S and M together. As in the case of Figure 4b, pairwise correlations were computed for single trials before any averaging. The histogram from this data for all the trials, irrespective of task type, for both monkeys, can be seen in Figure 4c. Once again, the correlations between the Observation, Reaction, Reach and Grasp phases were found to be significantly different as demonstrated by a Repeated Measures ANOVA (th(1.735,671.303) = 17.53,th < 0.001, Greenhouse-Geisser correction for sphericity). Post-hoc analyses showed that there was an increase in correlation upon the transition to the Reaction and Reach phases while the shift to the Grasp phase was accompanied by a decrease in mean absolute correlations (Holm’s test, th < 0.001 for all transitions).

### Cluster analysis on the LFP spectral power in the delta band

We then applied the cluster analysis to the LFP spectral power in the delta band. This was done in order to obtain a finer grained picture of correlations between the electrodes. We first make a qualitative report that demonstrates how the clustering is not restricted between the electrodes of one area. This is followed by a more quantitative analysis in which we follow the cluster number as a function of task epoch.

### Clustering of LFP activities between and within areas of the motor cortex

The spectrograms of Figure 5 were obtained by applying the cluster analysis to two trials of monkey M. Figure 5a and b come from two different sessions of monkey M. The clustering algorithm was applied to the entire trail. We can see in this figure that while electrodes from a same region often had similar activity (indicated by long patches of the same color in the left column coding for the region of origin of the electrodes), electrodes from different areas could also have correlated activity (indicated by the mix of color in this column). This highlights the idea that the reach-and-grasp movement is accompanied by inter as well as intra-regional correlations.

**Figure 5.**
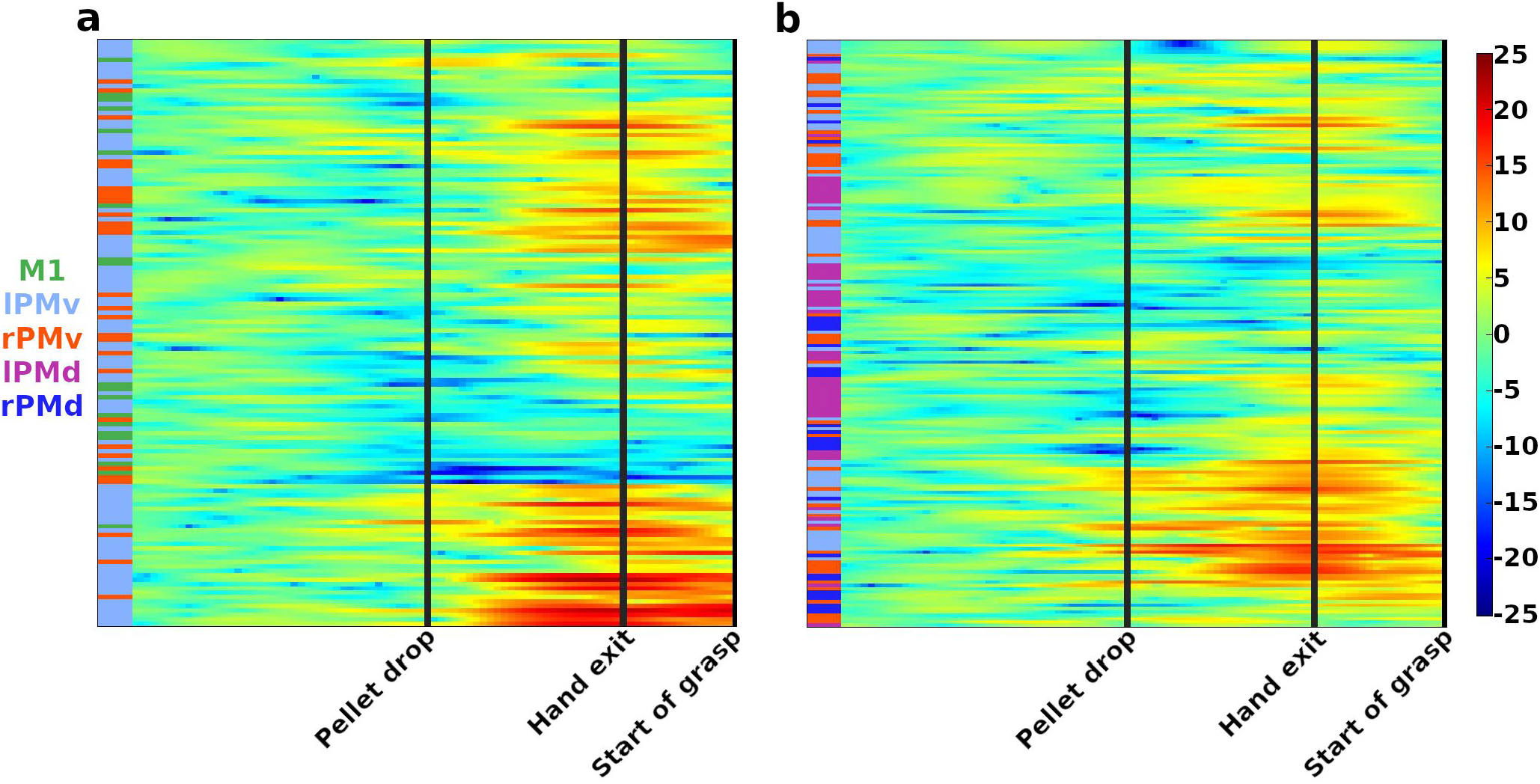
Clustering of the activity in delta band recorded from all the electrodes during one trial of monkey M. As in **Figure 3**, each line shows activity recorded from one electrode during the trial. The area of origin of the electrode is color coded by the column on the left of each graph (color legend on the left). It is to be noted that clusters are not restricted to a particular area of the motor cortex and contain LFPs from different regions. The black lines indicate the timing of the task epochs. a) and b) are from two different sessions of monkey M.

### Transitions in cluster numbers as a function of task epoch

Using absolute pairwise correlations of activity in the delta band provided us with a general idea of the similarities in the recorded LFP activities during the whole reach-and-grasp task. However, in keeping with the notion of synergies between muscles formed during a movement, we wanted to have a more detailed picture of grouping patterns in the LFP activity as the task progressed. We did this by using the hierarchical clustering method described in the Methods section. For this we increased the temporal precision of the analysis by using smaller temporal subsections for the clustering analysis. We divided each of the previously defined task epochs into 5 equal sub-periods and computed the cluster number in each sub-period. Figure 6 presents normalized cluster numbers (see Methods) as a function of task progression in these sub-periods.

**Figure 6.**
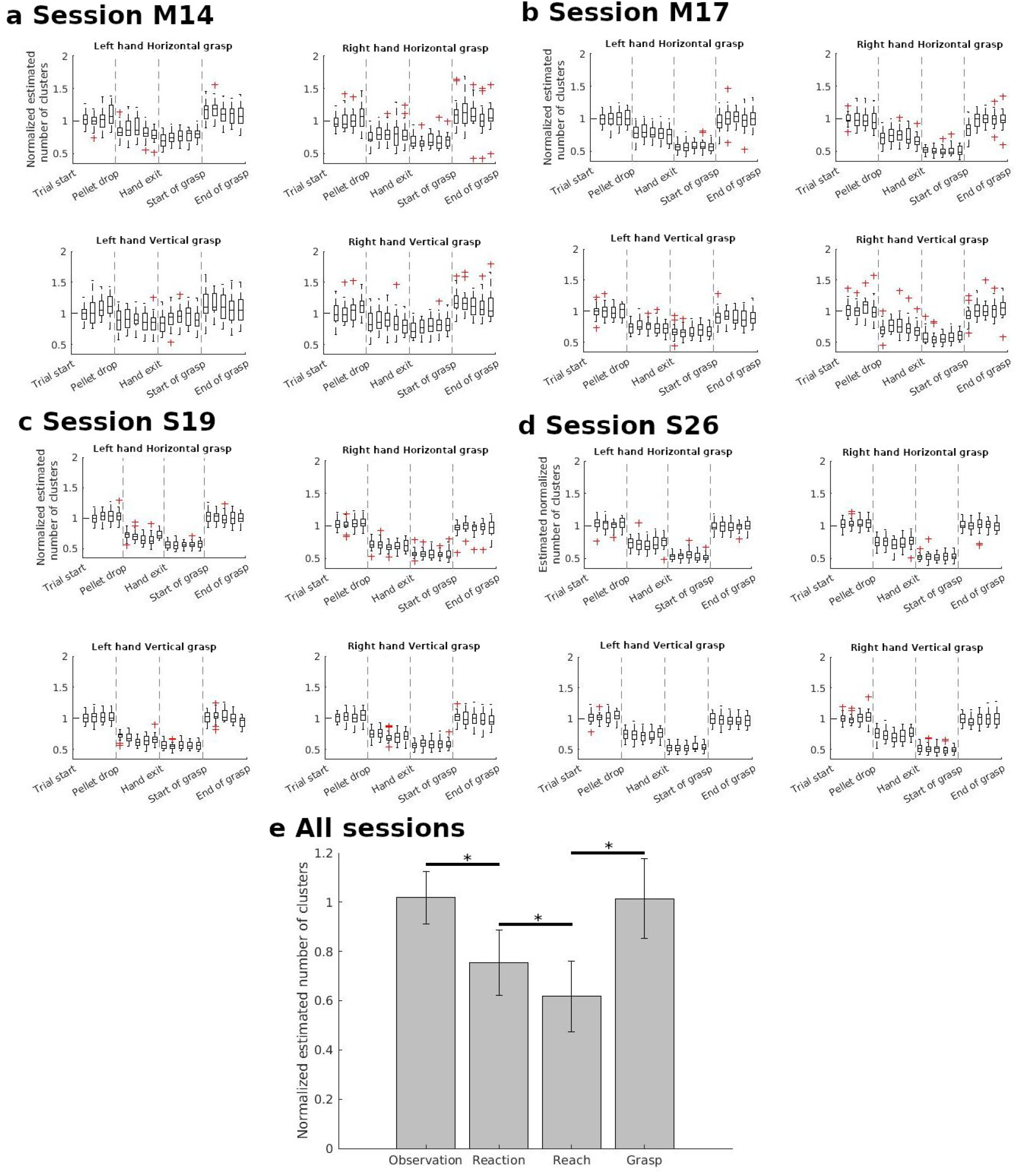
Average normalized cluster number in the delta band as a function of task epochs. a) and b) For all the trials of the same task conditions for monkey M. There are trials from two sessions presented. c) and d) For all the trials of the same task conditions for monkey S (over two sessions) e) For all the trials of both sessions of both monkeys. In this case, the average normalized cluster number for each task epoch is the average of the values from the five sub-periods. Error bars indicate the variance.

As in the case of correlation analyses, we first show the number of clusters of one session for each monkey (Figure 6a, b, c, d). Each figure displays the evolution in number of clusters as a function of trial epochs. The titles indicate the task conditions for each figure. They show that for both monkeys, independently of the arm used and grasp orientation, there was a relatively similar profile of clustering transitions. First, there was a decrease in the estimated number of clusters at pellet drop, followed by a smaller drop at Reach onset before going back close to baseline values during grasp. Such transitions in normalized cluster numbers are coherent with the results of the correlation analyses (Figure 2). Increased averaged pairwise correlation would indicate greater similarities across the activity of different electrodes which in turn would result in a smaller number of clusters.

To confirm the statistical significance of these transitions, we used the data obtained across both sessions of the two monkeys. Figure 6e displays the averaged normalized estimated cluster number during each of task epochs across all the sessions for both monkeys. Unlike Figure 6a-d, there is only one value of normalized cluster number for each task epoch, which is the average of the cluster numbers from the five sub-periods. A Repeated Measure ANOVA showed a significant effect of task epoch for cluster number ( *F*(2.751,1045.551) = 2795.862,*P* < 0.001, Greenhouse-Geisser correction for sphericity). Post-hoc analyses indicated significant differences in the normalized cluster number for each of the transitions in Figure 6e (Holm’s test, *p* < 0.001 for all the transitions).

### Prediction

The section above showed that cluster number evolves as a function of task epoch. The next step of our analysis was to examine if this variable had some promise for predicting changes in the phase of the movement ahead of time. To answer this question, the clustering was performed using a sliding window of 100 ms instead of the sub-periods used in the previous section. The sliding window is also more pertinent to an online analysis. The result of the sliding window analysis for one example trial is presented in Figure 7a.

**Figure 7.**
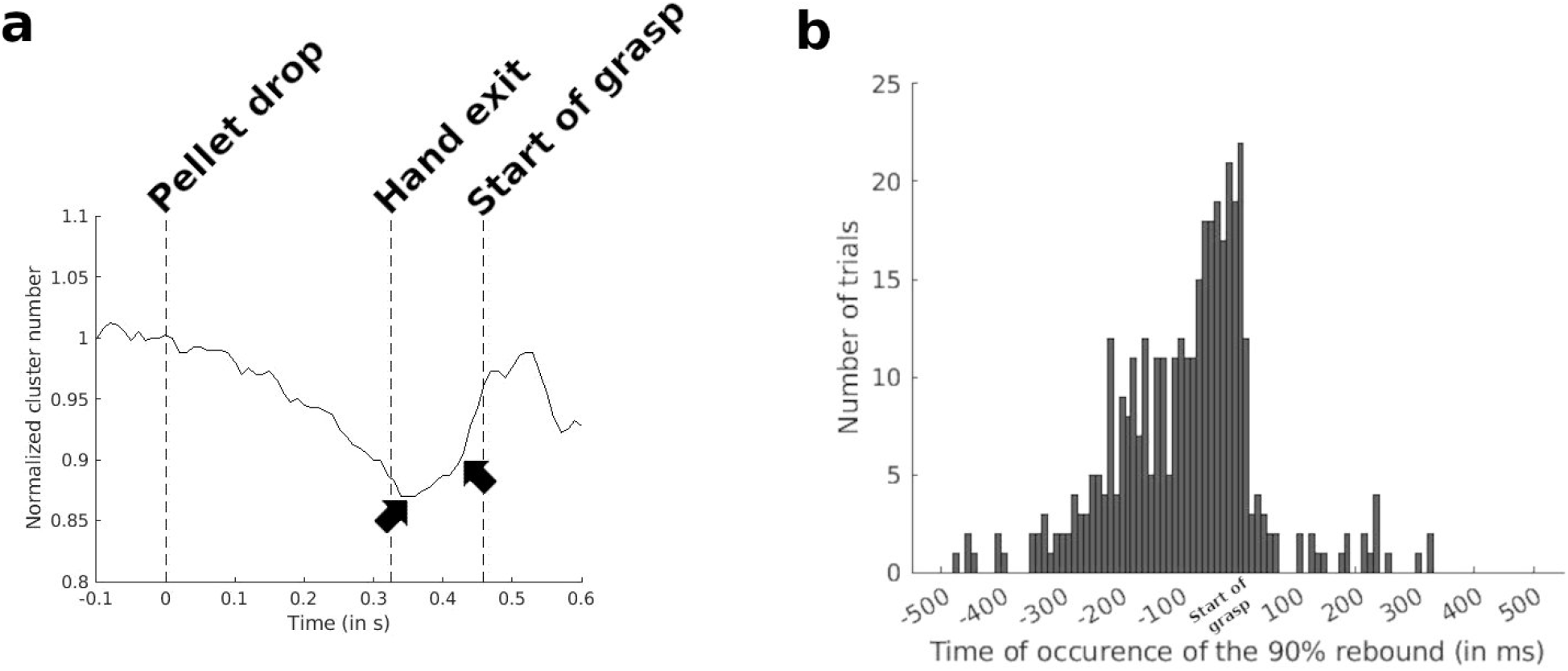
a) Smoothed trajectory of continuous estimated cluster number for one trial of a session of monkey S. The first black arrow indicates the local minimum between Pellet drop and Start of grasp. The second black arrow shows the rebound point where normalized cluster number reaches 0.9 and that the Start of grasp is imminent. From all recorded trials, a total of 18 trials (5.3%) did not show a minimum before Start of grasp. b) Distribution of the number of trials with time occurrence of the rebound to normalized cluster number 0.9 as a function of time delay from the Start of Grasp. All delays are aligned to the Start of Grasp as t=0. Negative values would indicate that the normalized cluster number reached 90% of baseline before the Start of grasp.

One particular feature of the continuous estimated cluster number was the occurrence in the vast majority of cases, of an inflection in the cluster number curve following pellet drop (Figure 7a). This feature resulted in the existence of a local minimum between Pellet drop and Start of grasp (first black arrow, Figure 7a). The occurrence of this minimum before start of grasp was observed for 94.7% of all recorded trials hence suggesting the possibility of using this feature of the LFP data to predict the Start of grasp. After this local minimum, the cluster number rebounded back in the direction of the baseline.

Despite being present for a vast majority of trials, the local minimum point was on average 250ms from the start of grasp. To find a point closer to the start of grasp, we decided it would be better to move beyond the minimum and up along the rebounding portions of the curve. To determine the optimal cluster number for providing a prediction point closer to the Start of grasp, we looked at the distribution of normalized cluster numbers at the Start of grasp for all the recorded trails. This distribution of normalized cluster number at Start of grasp showed a peak at 0.9 (90% of baseline), hence indicating that this would be an optimal threshold value with which to predict the imminent onset of Grasp. Figure 7b shows the result of this analysis. For all recorded trials we computed the distance from Start of grasp to the rebound point described previously. The distribution of these values showed that for most trials the rebound happens in a window of 80ms preceding the Start of grasp, hence indicating that using the rebound point of 0.9 provided a much closer prediction of the Start of Grasp. A chi-square test comparing the distribution of Figure 7b with that of a random distribution, showed a significant difference (Pearson’s chi-square test, χ^2^(80) = 599.674,*p* < 0.001). This confirmed the presence of a bias in the rebound values for certain intervals before the start of grasp.

## Discussion

In this study we focused on how grouping in LFP activities recorded by multiarray electrodes evolves as a function of a reach-and-grasp task. The analysis was done using correlations and hierarchical clustering. Correlations were high when Reach started and decreased by the time of grasping. The increase in correlations with reach is what would be expected from several previous studies at the behavioral and muscular level. Researchers have demonstrated a quasi-linear relationships between the kinematic angles of the arm during reach (Lacquaniti et al., 1986; Soechting & Lacquaniti, 1981). At the muscular level, indications of correlations during reach have also been demonstrated with the computation a small number of synergies which are able to reproduce the muscular activation patterns for reaching at different speeds, in different directions and with different loads (d’Avella et al., 2008, 2011; Muceli et al., 2010). Covariations in muscular activations patterns have also been described for finger movements (D’avella & Lacquaniti, 2013; Geed & van Kan, 2017; Klein Breteler et al., 2007; Pei et al., 2022; Weiss & Flanders, 2004). However, correlated activity during grasp can be expected to be lower than what is found during reaching as the fingers have to act in a more individuated manner. Previous research has shown that the number of components required to capture the variance during reach (d’Avella et al., 2008; Muceli et al., 2010) has been generally smaller than those needed for tasks with the hand (Mollazadeh et al., 2014; Takahashi et al., 2017; Weiss & Flanders, 2004). In keeping with this, the LFPs in our current study showed a decrease in correlation and increase in the cluster number as Grasp began.

Rather than working with mean values and relying on statistical effects, we chose to examine the patterns of correlations and cluster formations separately for each type of reach-and-grasp. This pattern of increasing correlation during Reach followed by a decrease during Grasp was very robust, following the same organization, irrespective of arm used and hand orientation. In future studies, it will be interesting carry out the same computation for other types of reach-and-grasp tasks to see if it is an invariant LFP characteristic for this type of motor task.

The algorithm chosen to investigate the grouping of the LFP activities in more detail was hierarchical clustering. Compared to PCAs, this method affords much greater ease of interpretation as each electrode belongs to one particular cluster as opposed to PCAs, in which the activity recorded by a given electrode can be distributed across several components. Hierarchical clustering was also chosen to avoid some difficulties found with other clustering algorithms. For example, with k-means (Lloyd, 1982) and k-nearest neighbors (Fix & Hodges, 1989) the number of clusters is determined a priori. Hierarchical clustering does not have this prerequisite. This method also provides a hierarchy of similarity between data points, hence, allowing for the progressive quantification of data similarity.

Our study joins several others that have found useful indicators concerning movement in LFP signals. Most of these did not investigate the question of correlations between LFP activities. For example, Mehring et al. (Mehring et al., 2003) used LFPs recorded within the motor cortex to infer which arm had been used as well as to decode hand position and movement velocity. The researchers found improved predictions by adding in the contributions of multiple electrodes but did not explore the question of movement information encoding in the correlations of LFP activities. Concerning movement planning, Scherberger et al. (Scherberger et al., 2005) showed how LFP activity before the start of reaching was different from that before a saccade in the posterior parietal cortex. This study used single electrode LFP recordings. Hence decoding concerning reach and saccades originated in single unit properties – in this case, the LFP amplitudes at specific frequencies. Spinks et al. (Spinks et al., 2008) demonstrated selectivity in the beta frequencies for six objects in the primary motor and ventral premotor cortex. They found poor correlation in the object tuning between the LFP recordings from M1 and the ventral premotor cortex of the monkey but did not seek to follow the nature of these correlations over the course of reach-and-grasp.

An example of a study focused on the question of correlations during reach-and-grasp is a study by Quarta et al. (Quarta et al., 2022). These researchers used hierarchical clustering to examine the evolutions of clustering in signals from calcium imaging during a reach-and-grasp task in mice. In agreement with our results, they also found a minimum Ward distance (which translates to a minimum cluster number in our study) right before Reach onset and a maximum at the Start of grasp. In contrast to our study, the Quarta et al. (Quarta et al., 2022) investigation with calcium imaging, was a more coarse grained study demonstrating how neuronal connectivity related to the reach-and-grasp could involve not only the motor cortex but several non-motor areas such as the visual, somatosensory and retrosplenial cortex. Our study with more than 100 electrodes in the motor areas provides a demonstration that a similar organization can be seen at a finer grained level.

Following the observations with mean values of cluster numbers during the sub periods of each epoch, we decided to conduct a sliding window analysis. Such a procedure would simulate an online monitoring of cluster numbers. Figure 7 shows how there is a progressive decrease in cluster number during the planning stage of reach which attains a minimum before a rebound of this value as grasp approaches. This local minimum could therefore serve as a marker of the imminent Start of grasp. We also explain in the Results section that following the rebound from the minimum point would take us closer to the actual onset of Grasp. Such a marker would be useful for prosthetic devices as it would indicate the point at which the LFPs have to be analyzed for the type of grasp that is to be performed. This would provide an easier classification task than one which analyzes the entire time series from the start of Reach. Other studies have reported on LFP features which can serve as temporal markers of task epochs. Both Scherberger et al. (Scherberger et al., 2005) and Hwang & Andersen (Hwang & Andersen, 2009) report that LFP power in the posterior parietal cortex increases for the low frequency bands and decrease for the high frequency bands during the phase between planning and execution for reaching. In contrast to the two previous studies in the posterior parietal cortex, an investigation by DePass et al. (DePass et al., 2022) was conducted in the motor cortex. They reported a better accuracy for distinguishing 5 task epochs during reach-and-grasp by using the LFP spectral power at high frequencies. They also report that in comparison, correlations between LFPs of the electrodes only provide a poor prediction of task epoch. We did not in our study attempt to distinguish 5 different task epochs but only the Start of grasp as a moment which would be key for the predictions of a prosthetic device. As close to 95% of the trials showed the global minimum in cluster number described in Figure 7, it indicates a high probably for good accuracies in the online automatic detection of imminent start of grasp. In future studies we will investigate different Machine Learning methods to provide these predictions in an automatic manner.

In our study the computation of correlations and clusters was carried out over all the implanted arrays of the motor cortex. This included the primary motor cortex for one hemisphere as well as the dorsal and ventral premotor cortexes of both hemispheres. This was in keeping with several human and animal studies that would seem to indicate overlapping activations of all these areas during the task. For example, there are observation from human studies of a close coordination between reach and grasp in which the two take place at the same time and influence each other (Jeannerod, 1984; Wallace et al., 1990). As a result of such observations at the behavioral level, Takahashi et al. (Takahashi et al., 2017) undertook an investigation in which they decided to investigate the classical separation of roles between the dorsal and ventral premotor cortex. They found that neurons in both area play a role in reach as well as grasp. Neural activity even in the case of unimanual movements is known to involve ipsilateral circuits as well (Moreau-Debord et al. 2021).

In conclusion, our study shows that alterations in correlations and cluster formation of LFP activity measured using multiarray electrodes in the motor cortex, are consistent with what would be expected from several lines of research at the behavioral level of the reach-and-grasp task. Hierarchical cluster analyses of LFP spectral power in the delta band showed a consistent decrease in normalized cluster number as Reach started and increased during Grasp. Since LFP recordings are easier and more robust to obtain than neuronal recordings, this shows a promise for the monitoring of motor control for muscular disorders involving abnormal synergistic activities

